# Crop modeling suggests limited transpiration would increase yield of sorghum across drought-prone regions of the United States

**DOI:** 10.1101/2023.06.27.546776

**Authors:** Rubí Raymundo, Greg Mclean, Sarah Sexton-Bowser, Geoffrey Morris

## Abstract

Breeding sorghum for drought adaptation is pivotal to secure crop production in drought-prone regions. Limited transpiration (LT) restricts water demand at high vapor pressure deficit, saving water for use in critical periods later in the growing season. Here we evaluated the hypothesis that LT would increase sorghum grain yield in the United States. We used a process-based crop model, APSIM, which simulates interactions of genotype, environment, and management (G × E × M). In this study, the G component includes the LT trait (G_T_) and maturity group (G_M_), the E component entails water deficit patterns, and the M component represents different planting dates. Simulations were conducted over 33 years (1986-2018) for representative locations across the US sorghum belt (Kansas, Texas, and Colorado) for three planting dates and maturity groups. The interaction of G_T_ x E indicated a higher impact of LT sorghum on grain for LD, MD, and ED (8%), than on WW environments (4%). Thus significant impacts of LT can be achieved in western regions of the sorghum belt. Otherwise, the lack of interaction of G_T_ × G_M_ × M suggested that an LT sorghum would increase yield by around 8% across maturity groups and planting dates. Although the interaction G_M_ × M revealed that specific combinations are better suited across geographical regions. Overall, the findings suggest that breeding for LT would increase sorghum yield in the drought-prone areas of the US without tradeoffs.

## INTRODUCTION

Droughts resulting from changes in precipitation patterns threaten crop production and food security in semiarid areas worldwide (Barbier, 2015). In the United States alone, crop yield loss due to droughts costs ∼$9 billion per year (NOAA, 2020). In this respect, breeding for drought-prone environments plays a pivotal role in maintaining crop production (Thornton et al., 2018). Nevertheless, developing crops with less water demand is challenging because drought adaptation traits are complex, difficult to identify, and often involve tradeoffs (Araus et al., 2012; Monneveux et al., 2012). Furthermore, testing the effect of these traits under water stress scenarios is limited since drought events vary over time and geographies (Pournasiri-Poshtiri et al., 2018; Tang and Piechota, 2009). Thus, plant breeding programs require complementary methods to test the effect of any hypothetical drought adaptation trait to design a breeding pipeline (Bernardo, 2020; Cooper et al., 2002; Cooper and Messina, 2022).

Crop models have become standard tools to assess the impact of new technologies in agriculture and can support plant breeding (Challinor et al., 2018; van Ittersum et al., 2003). These models integrate ecophysiological knowledge to represent the plant-soil-atmosphere system and predict the crop response to soil properties, climatic conditions and crop management practices (Jones et al., 2003). Crop models equip breeding programs with the tools to develop and evaluate hypotheses regarding the performance of new cultivars (G) under environmental (E), and management scenarios (M) (Chenu et al., 2017; Messina et al., 2011). Several crop modeling studies have evaluated theoretical expressions of crop traits linked to cultivar-specific parameters for drought environment (Singh et al., 2014). The most common approach varies cultivar parameters (Messina et al., 2011; Singh et al., 2014) or implements new traits (Sinclair et al., 2005) to evaluate alternative ideotypes for constraint environments. This approach to crop improvement advantages investment of finite resources to defined targets for genetic gain in specific environments.

Sorghum is one of the most drought-adapted crops in semiarid regions used for multiple purposes, including forage, fiber, and food (Doggett and Majisu, 1968; Smith and Frederiksen, 2000). Most of the grain sorghum production worldwide (15%) is grown under rainfed environments in the sorghum belt of the United States that runs from South Dakota to South Texas (Laingen, 2015). Kansas, Texas, and Colorado lead grain sorghum production in the sorghum belt with 50%, 30%, and 6%, respectively (Laingen, 2015). Across this area, water limitation and high vapor pressure deficit (VPD) affect plant transpiration, making sorghum production vulnerable to droughts. Although sorghum harbors drought adaptation (Abdel-Ghany et al., 2020; Abreha et al., 2021), breeding for drought traits has received less attention. Therefore, the full potential of sorghum production under water-limited environments in the sorghum belt of the United States may not yet have been achieved.

Limited transpiration (LT) is a hypothetical trait that restricts water demand in periods of high VPD which occurs around mid-day (Figure 1A and 1B). This mechanism shifts plant-water demand, conserving water in the soil profile during the vegetative stage and for use during grain filling (Figure 1) (Sinclair et al., 2005). Reducing transpiration (H_2_O) due to stomatal closure in hours with high VPD would penalize carbon assimilation (CO_2_). Thus, causing grain yield reductions under well-watered conditions but increasing the grain yield and the effective use of water under moderate water-limited environments (Vadez et al., 2014). This hypothetical physiological mechanism of LT has been extended into process-based models where transpiration was restricted during high VPD hours (Messina et al., 2015; Sinclair et al., 2005; Truong et al., 2017). Crop model simulations under rainfed conditions for sorghum and other crops such as soybean, maize, lentil, chickpea, and wheat indicate a yield increase for a phenotype with LT trait in areas vulnerable to water scarcity (Collins et al., 2021; Sinclair et al., 2017). For sorghum, reports indicated an increase in yield production from 6 to 10% for severe drought scenarios in Australia, semiarid regions of India, and the United States (Texas) (Kholová et al., 2014; Sinclair et al., 2005; Truong et al., 2017).

**Figure 1.**
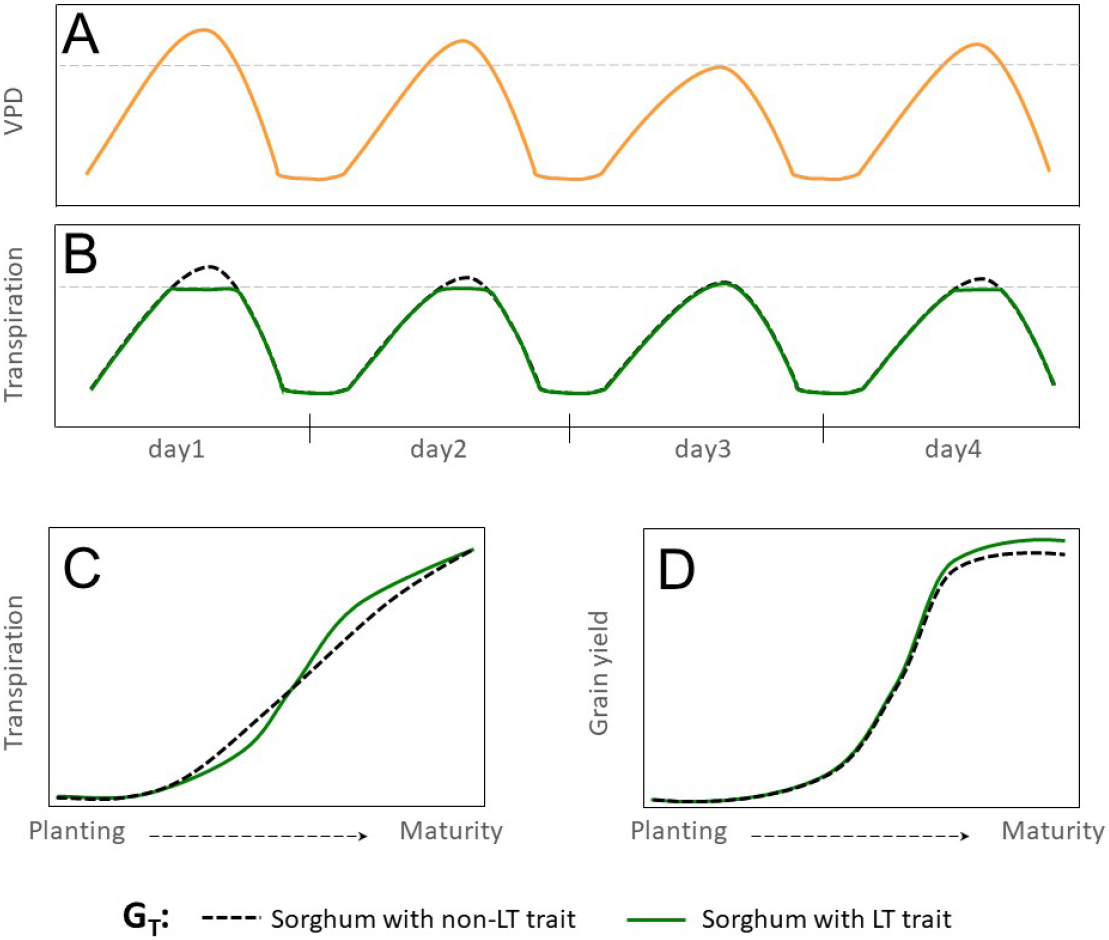
Hypothetical effect of LT trait in grain sorghum under rainfed conditions. (A) Hourly trajectory of VPD during four days with contrasting, (B) Hourly trajectory of transpiration for sorghum with non-LT and LT traits (G_T_). The dashed gray line indicates the threshold of VPD at which genotypes express differences in transpiration VPD response. (C) Cumulative transpiration during the growing season, and (D) trajectory of grain yield during the growing season.

Simulations for various crops (Collins et al., 2021; Guiguitant et al., 2017; Messina et al., 2015) suggest that breeding for the LT trait can make a valuable economic contribution in rainfed regions. Yet, its impact on grain yield in sorghum-producing regions of the United States remains unknown. This study uses the APSIM-sorghum growth model to generate hypotheses of the potential benefits and tradeoffs of the LT trait in grain sorghum. Under drought scenarios, we expect an increase in grain yield in rainfed sorghum-producing regions for sorghum with the LT trait (Figure 1). Otherwise, no impacts or detrimental effects on grain yield are expected for non-stress environments. Likewise, we expect these benefits across different combinations of genetic background and management practices. Results indicate that introgressing LT in grain sorghum would increase yield by more than 5% in water-limited scenarios but less than 5% in well-watered settings. Additionally, the LT would benefit grain yield across all combinations of maturity groups and planting dates.

## MATERIALS AND METHODS

### Production system and study sites

The simulation study was conducted for Kansas, Texas, and Colorado counties that have high sorghum production (Figure 2A) area and are located in contrasting gradients of precipitation and VPD (Table 1, Figure 2D). Across these locations annual precipitation and VPD are inversely associated (Figure 1B, 1C, and 1E) with declining precipitation and increasing VPD from east to west (https://prism.oregonstate.edu/). Annual precipitation shapes farmer crop management including maturity group adoption (Ciampitti et al., 2019; Roozeboom and Fjell, 1998; Shroyer et al., 1998). Therefore, in these regions plant density of 17 plants m^2^ and 6 plants m^2^ are recommended for areas with annual precipitation around 800 mm and 350 mm, respectively (Shroyer et al., 1998). Similarly, full-season hybrids are planted in regions with high annual precipitation while short-season hybrids are grown in regions with low precipitation (Ciampitti et al., 2019; Roozeboom and Fjell, 1998).

**Table 1.**
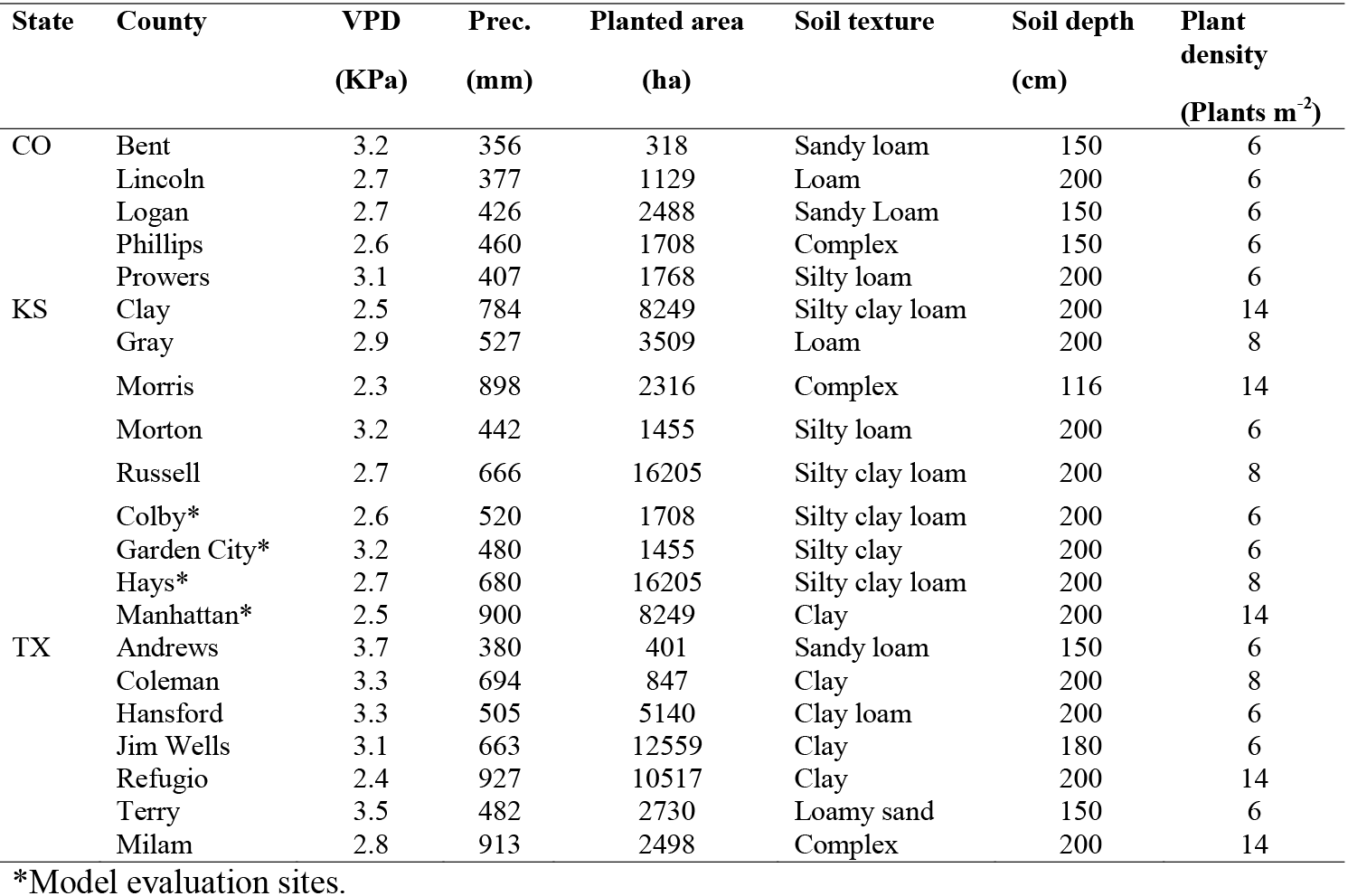
Characteristics for the study locations across the US sorghum belt.

**Figure 2.**
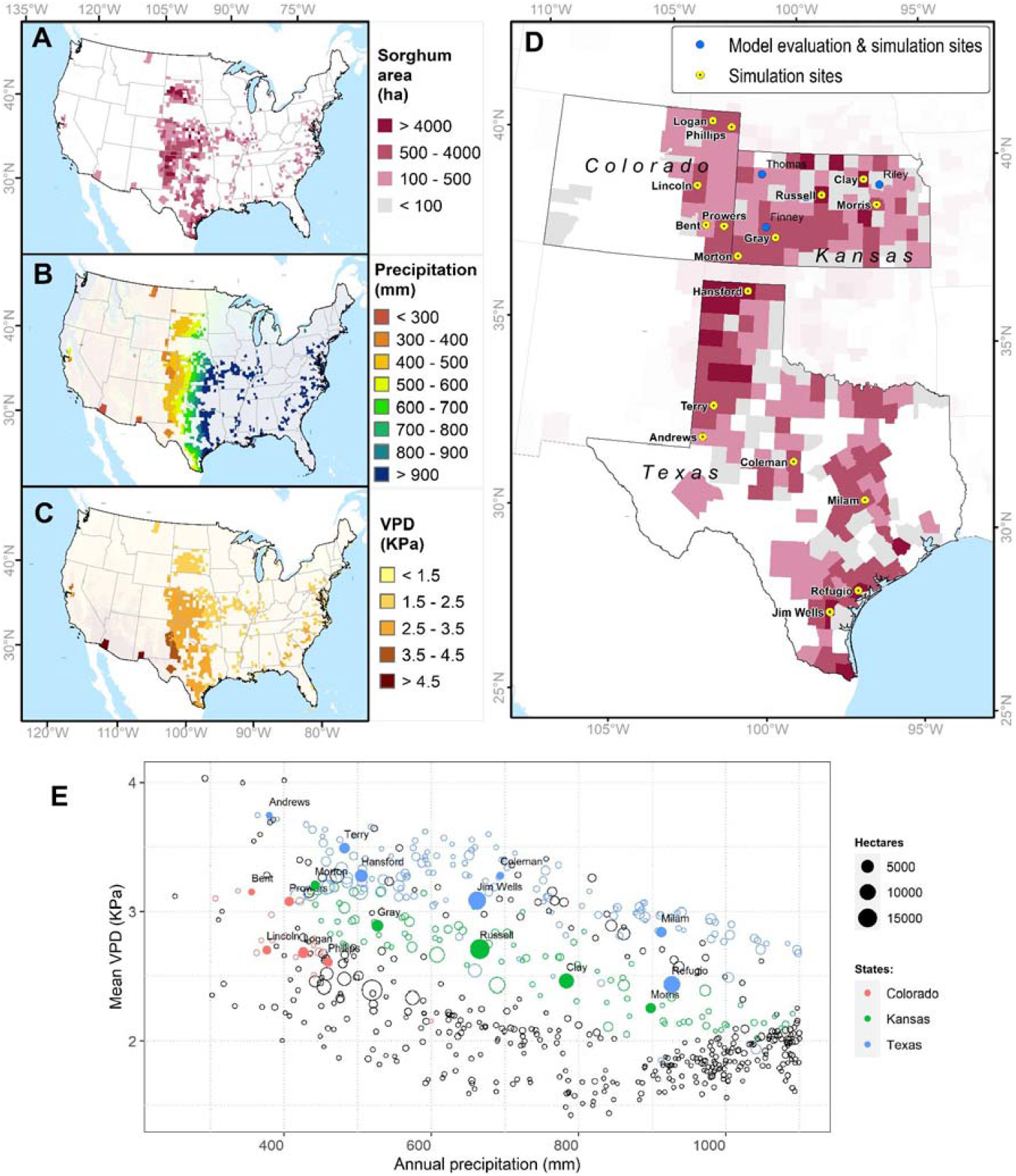
Study system to evaluate the impact of the limited transpiration (LT) trait. (A) Geographic distribution of grain sorghum production area (ha^-1^) in the United States in 2019. (B) Annual precipitation (mm) in sorghum producing areas. (C) Mean of the monthly maximum vapor pressure deficit (VPD, kPa) from May to August in sorghum producing areas. (D) Distribution of grain sorghum in Colorado, Kansas, and Texas and sites for model evaluation and simulation sites. (E) Mean VPD and annual precipitation in sorghum producing regions. Sorghum production areas were obtained from the National Agricultural Statistical Service (NASS, https://www.nass.usda.gov/). Precipitation and vapor pressure deficit information for sorghum-producing areas were acquired from the PRISM Climate Group (https://prism.oregonstate.edu/).

### APSIM-sorghum crop model

APSIM-Sorghum (Hammer et al., 2019, 2010) is a crop model that integrates the intertwined interaction of G × E × M) to simulate plant development and growth on a daily basis (Holzworth et al., 2014; Keating et al., 2003; Wang et al., 2002). The model requires the following input data: daily weather records, soil profile characteristics, crop management, and cultivar-specific parameters. Crop phenology development is estimated as the summation of thermal time for nine phases from germination to physiological maturity. Daily biomass is estimated as the minimum of biomass limited by solar radiation or water availability. The biomass limited by solar radiation is the product of radiation use efficiency (RUE, MJ m^2^), solar radiation (MJ m^2^), and the fraction of light intercepted. The biomass limited by water availability is the product of transpiration efficiency and soil water supply. The model estimates water, temperature, and nitrogen deficit factors which affect phenology and growth. To estimate the effect of LT on carbon assimilation in hours with high VPD, APSIM-sorghum downscales daily temperature and solar radiation to hourly time steps and estimates relative humidity (RH) on each hour (Parton and Logan, 1981). Temperature and RH are used to calculate VPD on each hour (Monteith and Unsworth, 2013; Murray, 1967), then the model estimates biomass as a function of hourly water supply and demand. Finally, the biomass accumulation is aggregated for each daily timestep. Note, the version of APSIM-sorghum used and LT modifications were made for research purposes and are not in the release version.

### Model inputs

Daily weather data at each site, including precipitation (mm), solar radiation (MJ m^-2^ day^-1^), maximum (°C), and minimum temperature (°C), were obtained from NASA Prediction of Worldwide Energy Resources (POWER-https://power.larc.nasa.gov/) from 1986 to 2018. The spatial resolution of the data are 1.0° latitude by 1.0° longitude for solar radiation and 0.5° latitude by 0.5° longitude for the remaining variables. Soil profile information such as soil texture (%), bulk density (g ml^-1^), organic carbon (%), and pH was downloaded from the web soil survey (https://websoilsurvey.sc.egov.usda.gov/App/HomePage.htm). These data were used to estimate the saturation capacity (SAT), field capacity (DUL), and wilting point (LL15) for each layer of the soil profile using the SBuild application of the Decision Support System for Agrotechnology Transfer (DSSAT) program (Hoogenboom et al., 2019). Crop management practices such as planting depth, plant population and planting dates were obtained from experiments or variety trials (Larson et al., 2021; Pachta, 2007; Schnell et al., 2021).

### Model testing

Model testing was conducted in two steps: model calibration and model evaluation (Wallach et al., 2014). In model calibration, specific parameters were iteratively adjusted to fit observations, while model evaluation estimated the accuracy of the model with independent data. For model testing we collected available information on field experiments for hybrid 87G57 from 1997 to 2007 (Figure 1D, Table S1). Model calibration was conducted with a high quality experiment that accounts for information of crop management, phenology development, in-season biomass components, and initial soil water (Pachta, 2007). Information of this experiment including weather, soil and crop management was arranged into APSIM-Sorghum. First, a simulation was conducted for the hybrid 86G56 (no calibration), which was available in the library of the model. Next, cultivar parameters were modified, to eliminate the photoperiod sensitivity (*photoperiod_slope* from 10 to 0), and to match the grain yield components by modifying the parameter relation between biomass accumulated from floral initiation to the start of grain (*dm_per_seed* from 0.00087 to 0.00099). There was no need to adjust parameters related to phenology development since the model was accurate in predicting flowering time for this experiment (observed: 52, and simulated: 53).

Model evaluation was conducted with variety trial experiments conducted in Garden City, Colby, and Hays (Kansas). These experiments have information of planting date, plant density, flowering time, and grain yield. Environment (daily weather data, soil profile) and crop management practices for these simulations were arranged into APSIM-Sorghum. Each year the crop was simulated to be planted under optimal soil moisture (70% soil available water), and non-nitrogen limitations at plant density of 6–14 plants m^-2^. Grain yield was expressed assuming 12.5% of moisture content. Model accuracy was analyzed using the root mean square error (RMSE), which indicates the distance from a perfect prediction (Wallach et al., 2014).

### Model application

Parameters for hybrid 87G57 corresponded to a commercial short-season sorghum hybrid with 15 leaves and a non-LT trait. The number of tillers was kept constant for all phenotypes. Parameters controlling growth and development, *tt_endjuv_to_init, Tpla_prod_coef*, and *Tpla_inflection*, were adjusted to simulate mid-season and late-season sorghum hybrids, each with 17 and 19 leaves, respectively. The number of tillers was kept constant (0.3) for all maturity groups. The model defines a phenotype with an LT trait by assigning the parameter *limited maximum transpiration* to any value from 0.2 to 0.9 mm h^-1^ (Table S1). Note a phenotype with a LT trait of 0.9 mm h^-1^ represents a genotype that restricts the transpiration by 10%. For simulations across all locations, the LT trait was imposed as 0.9 mm h^-1^. Simulations for sorghum with LT and non-LT traits started every year on the first of January with initial soil moisture of 60%. In these simulations, the crop was automatically planted at three time intervals, early-May, mid-May, and early-June, a row distance of 76 cm, planting depth of two cm, and fertilized to fully meet plant nitrogen demand. Simulations were conducted every year from 1986 to 2018. We analyzed the grain yield, transpiration, and soil moisture for both sorghum phenotypes (non-LT and LT).

We conducted a sensitivity analysis in a representative location to identify initial conditions’ effect on grain yield changes resulting from the LT trait. Therefore, simulations started with initial soil moisture of 20%, 30%, 40%, 50%, 60%, 70%, 80%, and 90% while maintaining the LT trait at 0.9 mm h^-1^. Otherwise, to determine yield gains resulting from hypothetical genetic variability, we created simulations and varied the LT parameter from 0.2 to 0.9 mm h^-1^ while maintaining the initial soil moisture at 60%. As previously outlined, these simulations started each year on the first day of January under similar maturity groups and management practices. Absolute and relative change in harvested grain yield for the phenotype with LT trait was calculated for each simulation and averaged over environments.

### Statistical analysis and interpretation

Statistical analyses of model outputs were performed in the R statistical environment utilizing mixed linear models and the *lmer* library (Bates et al., 2015). The analysis quantified the size of fixed effects on grain yield. Factors with fixed effects were trait (G_T_), maturity group (G_M_), planting date (M), water stress environments (E), and their interaction while factors with random effects were years nested on each location.

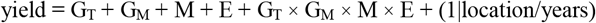

Tukey’s test was performed when the *F* value was below an α < 0.05 significance threshold. In this study, the water stress environment (E) was determined for non-LT sorghum which resulted from seasonal trajectory of daily water stress simulated for each season. This time series information was analyzed via a hierarchical classification using the *cluster* (Maechler et al., 2022) library and the *clara* (Clustering Large Applications clara) method (Kaufman and Rousseeuw, 1990). The number of clusters was determined via the silhouette method (Kassambara, 2017), a measure of similarity for each data point relative to the assigned cluster and other clusters. The final water stress patterns resulted as the median of water stress on each cluster.

## RESULTS

### Accuracy of model for grain yield prediction

To determine the model accuracy for flowering time and grain yield, we compared the observed data versus the information simulated by the model. For a growing season with hybrid 87G67 in Manhattan, Kansas (Figure 3A, 3B, 3C, and 3D), the model reproduced the trajectory of dry weight for total biomass, stem, and panicle with an RMSE of 1.1, 0.4, and 0.7 Mg ha^-1^, respectively. However, a substantial underestimation occurred for dry leaf weight. In this experiment, the observed grain yield was 4.8 Mg ha^-1^, and the results after calibration were 5.4 Mg ha^-1^. For experiments in Kansas from 1997 to 2007 in Garden City, Hays, Colby, and Manhattan, the model showed satisfactory predictions for days to anthesis with an RMSE of 5 days (Figure 3E) and grain yield with an RMSE of 2 Mg ha^-1^ (Figure 3F). Despite the lack of experimental field data for model testing in Texas and Colorado, a comparison of statistical (2.2 to 6.6 Mg ha^-1^) versus simulated grain yield (1.5 to 6 Mg ha^-1^) resulted in a RMSE of 1.1 Mg ha^-1^ (Figure 3G).

**Figure 3.**
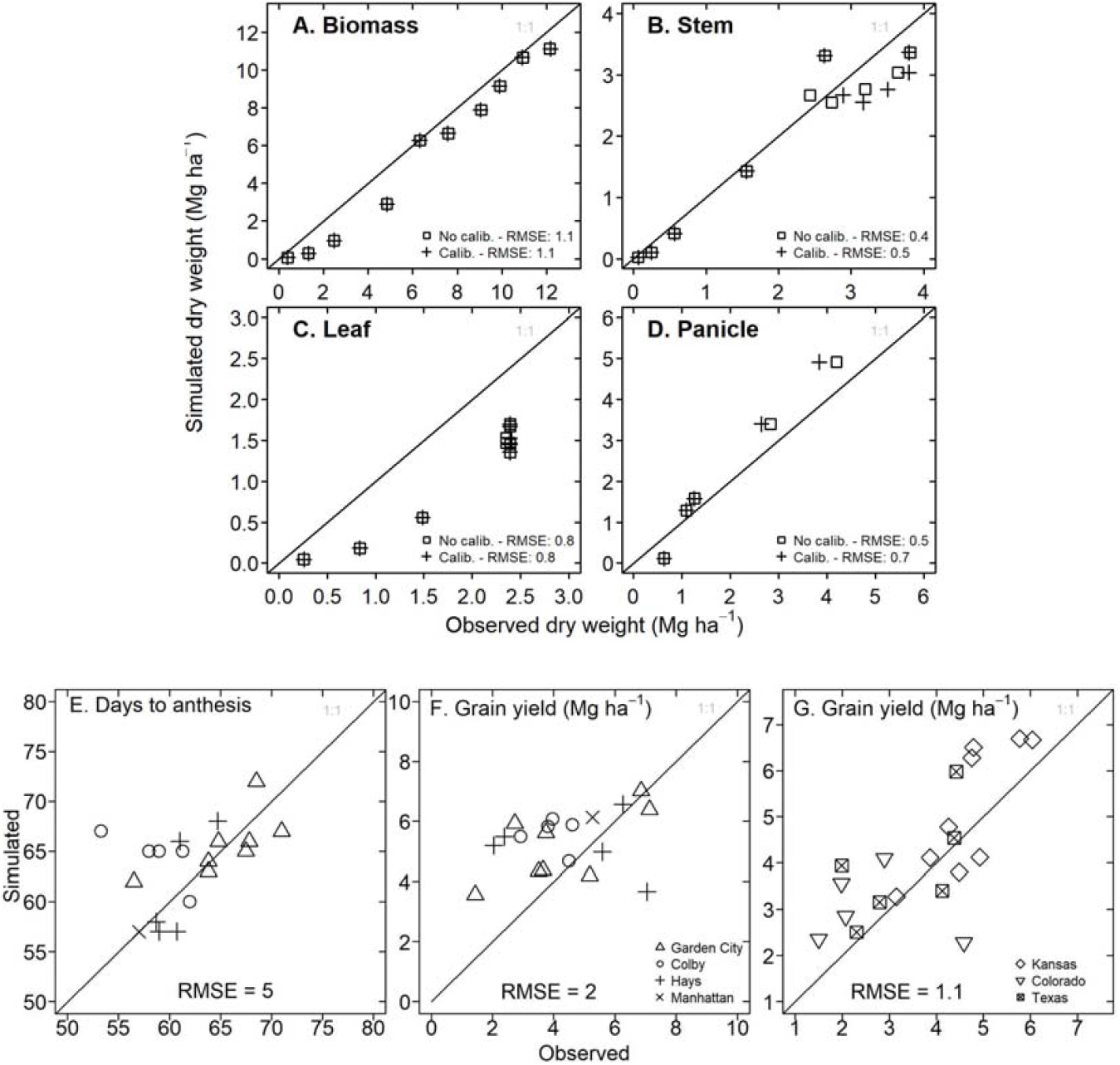
Model testing for APSIM-Sorghum in the study system. (A) Observed versus simulated in-season dry biomass for hybrid 87G67 in Manhattan, Kansas (2007). (B) Observed versus simulated in-season stem dry weight for hybrid 87G67 in Manhattan, Kansas (2007). (C) Observed versus simulated in-season leaf dry weight for hybrid 87G67 in Manhattan, Kansas (2007). (D) Observed versus simulated in-season panicle dry weight for hybrid 87G67 in Manhattan, Kansas (2007). (E) Observed versus simulated days to anthesis for hybrid 87G67 under rainfed conditions for experiments in Garden City, Colby, Hays, and Manhattan (Kansas). (F) Observed versus simulated dry grain yield for hybrid 87G67 under rainfed conditions across the Kansas precipitation gradient (Garden City, Colby, Hays, and Manhattan). Each point (Figures E and F) represents results for single seasons between 1997 to 2007. Information of (G) Observed versus simulated grain yield for Kansas, Colorado, and Texas study sites (indicated in Figure 4D). The *y* axis represents the mean of simulated yields over 33 years (1986-2018), three planting dates, and three maturity groups. The *x* axis represents the mean of observed data over 21 years (2010 to 2021). Observed sorghum grain yield (Figure G) was obtained from the National Agricultural Statistical Service (NASS, https://www.nass.usda.gov/).

### Variation of grain yield across G_M_ × M scenarios in the absence of LT

To determine the best G_M_ and M combination for grain yield in precipitation gradients, we conducted simulations for short-, medium- and full-season sorghums planted in early May (May 01), mid May (May 15) and early June (June 15). Note, around 3% of the simulations were removed for the analysis because they did not complete the vegetative stage (hereafter referred to as “failed seasons”) and the yield was close to zero. This occurred under extreme drought events (Rippey, 2015). For instance, in Colorado in 2012, the annual precipitation was less than 207 mm, and the rainfall during the simulated growing period was less than 130 mm. The number of failed seasons for full-season sorghum either planted early or late was higher in Colorado suggesting that short season varieties outperform any maturity group under low rainfall, while the frequency of failed seasons in Texas was higher in early planting dates (Figure S2).

Grain yield for simulated sorghum with a non-LT trait for different maturity groups and planting dates in Kansas, Texas, and Colorado, are indicated in Figure 4. An average across G_M_ and M indicated that grain yield varied from 1.7 to 6.5 Mg ha^-1^ (Figure 4A), with higher and lower yields in eastern and western regions, respectively. Interannual variability for grain yield ranged from 30 to 50% (Figure 4A). The model predicted higher yields in Colorado and Kansas when all maturity groups were planted in early May, followed by planting dates in mid-May and early June. In most Texas locations, the model predicted a higher yield for planting dates in early June (Figure 4B). The seasonal rainfall during each simulated season influenced the performance of different maturity groups for grain yield (Figure S1A). On planting dates in June, discrepancies among maturity groups occurred under high precipitation; nevertheless, as the amount of rainfall during the growing season decreases, these differences become negligible (< 1%; Figure S1B). By contrast, differences in maturity groups for grain yield across precipitation gradients in early May are insignificant (*p* > 0.05) (Figure S1B).

**Figure 4.**
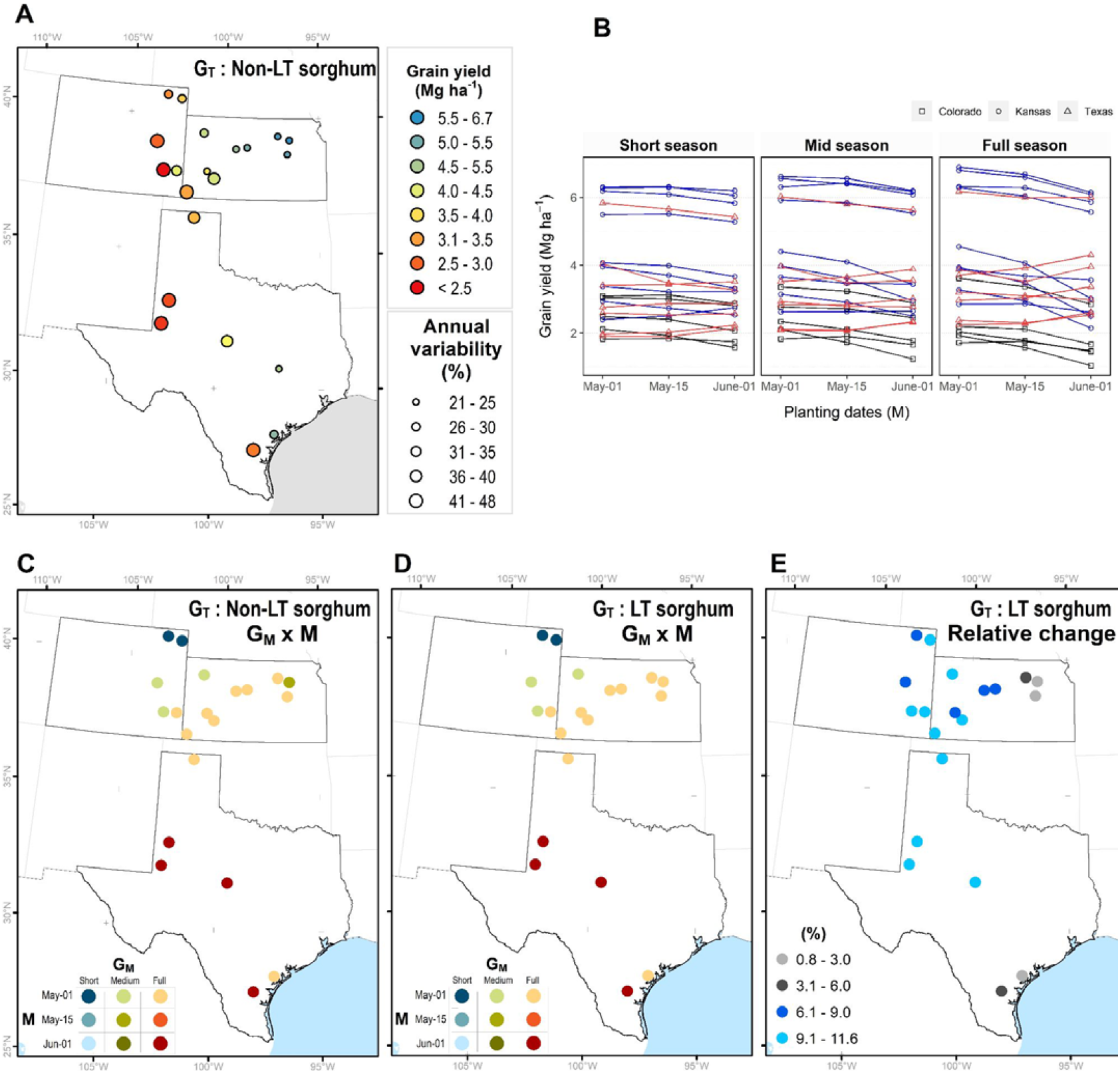
Maturity (G_M_) × planting date (M) combinations to achieve maximum yield for non-LT versus LT sorghum. (A) Average grain yield and interannual variability. (B) Average grain under different planting dates and maturity groups. Each point represents the mean of 33 years (1986-2018), the annual variability (coefficient of variability) is the quotient of the standard deviation and mean. Best G_M_ × M combination for a sorghum with (A) non-LT and (B) LT trait. Relative increase in grain yield for a sorghum with LT trait.

### Effect of LT on the best combination of maturity group and planting date across sites

Due to G × M × E interactions, the effect of non-LT vs. LT trait (G_T_) may depend on agronomic options, such as maturity group (G_M_) of the hybrid and planting date (M). To identify the best combination (G_M_ × M) at each site (E), we obtained the maximum yield for LT sorghum. The model indicated similar combinations for non-LT and LT sorghums (Figure 4C–D). In Colorado and Kansas, higher yields resulted when seasons for all maturity groups started on the first of May. In Colorado, short-season sorghum performed better in northern regions, while medium- season sorghum in southern regions. Full-season sorghum yielded higher across Kansas, except in Colby, where medium-season sorghum outperformed any other combination. In Texas, the model indicated full-season sorghum planted on the first of June as the best combination, with some exceptions in the northern regions (i.e. Hansford). Overall, sorghum with LT across all locations is expected to increase grain yields from 0 to 15% (Figure 4E). Note, the relative change is lower than 3% in regions with high precipitation and this change increases as declining precipitation amplifies water deficit scenarios in western regions of the study site.

### Water deficit environments are more recurrent in the West

To evaluate the value of LT in target population environments, we determined water stress patterns by clustering simulated time series information on water supply and demand (waterSD). The classification indicated four water deficit environments: well-watered or light stress at grain filling (WW), late drought (LD), mid-season drought (MD), and early drought (ED) (Figure 5A). WW and LD predominated in eastern regions of Kansas and Texas, while MD and ED predominated in eastern Colorado and western Texas (Figure 5B). The analysis revealed a strong correlation between seasonal precipitation and water deficit patterns; although it was non- significant (*r* = 0.9, *p* < 0.06). On average, WW, LD, MD, and ED, showed seasonal precipitation of 400 mm, 300 mm, 244 mm, and 230 mm, respectively.

**Figure 5.**
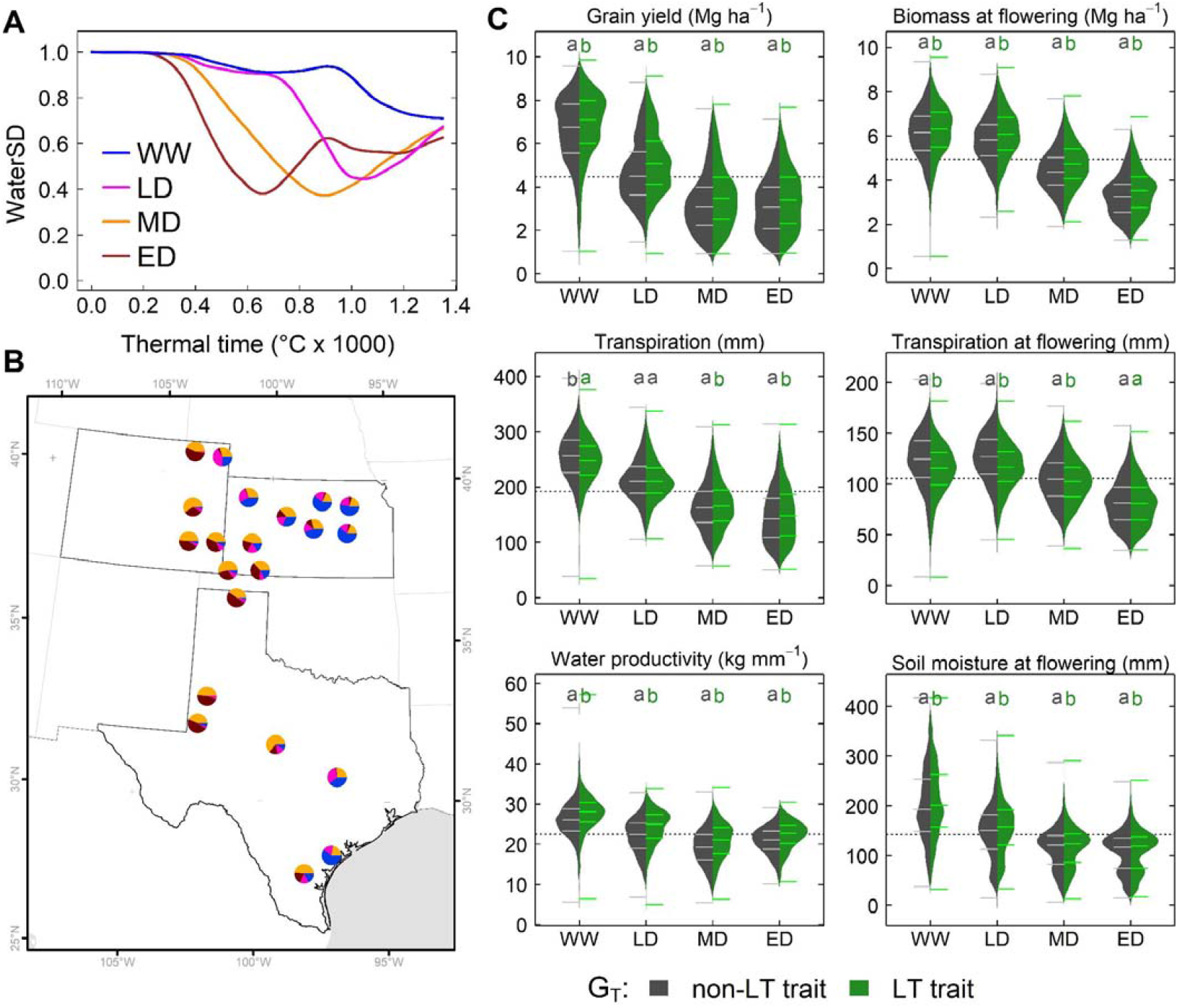
Simulated effects of LT (G_T_) across the US sorghum belt. (A) Water stress environment identified via water supply and demand (WaterSD) at each site. (B) Frequency of water stress environments in Kansas, Texas and Colorado. (C) Distribution of simulated grain yield, transpiration, water productivity, biomass and soil extractable water for a phenotype with a non-LT (darkgray) and LT trait (green) in water stress environments. Each distribution represents simulations for all maturity groups (G_M_), and planting dates (M) in all sites over 33 years. Letters indicate significant differences (α < 0.05) of all pairwise comparisons using the Tukey HSD test.

### The effects of L G_T_ × G_M_ × M × E

To determine significant interaction of G_T_ × G_M_ × M × E, we used a mixed model to compare variances across the mean (Table S3). We conducted this analysis for dependent variables at the end of the growing season (grain yield, transpiration and water productivity) and flowering time (biomass, transpiration, and soil water content). All individual fixed effects (G_T_, G_M_, E, and M) had high significance (*p* < 1×10^−4^, Table S3), and a post hoc analysis suggested that factors on each fixed effect belonged to different groups (Table S3). For instance, sorghum’s LT trait increased grain yield by 5%. Full-season sorghum yielded 21% and 10% higher than early and mid-season sorghum. WW favored grain production; while lower yields correspond to ED. Likewise, earlier planting dates outperformed sorghum planted either in mid-May or early-June.

All dependent variables exhibited high significance (*p <* 1×10^−5^) in two-way interaction for three combinations: G_T_ × E, E × M, and G_M_ × E (Table S3). The significant interaction for G_T_ × E, and the pairwise comparison for grain yield (Figure 5C), water productivity (Figure 5D), biomass at flowering (Figure 5C), and soil moisture at flowering (Figure 5H) indicated that the LT trait outperformed the non-LT trait in all environments. Note, grain yield for a LT sorghum in a WW environment was lower (4%) than in LD, MD, and ED environments (8%). However, the pairwise comparison for total transpiration and transpiration at flowering confirmed the significance for the interaction G_T_ × E. For instance, the non-LT trait exhibited higher total transpiration in WW, while the LT trait improved it in MD and ED (Figure 5C). At flowering time, LT increased transpiration in WW, LD, and MD, but both traits exhibited similar transpiration in ED (Figure 5C). Only for biomass at flowering time and water productivity the interaction G_T_ × M had high significance.

Transpiration at flowering exhibited a three-way interaction for G_T_ × G_M_ × E (*p <* 0.01). The pairwise comparison indicated a lack of differences between LT and non-LT genotypes for each maturity group in WW (Figure S4A, S4B, and S4C). By contrast, the LT trait increased transpiration in WW, LD, and MD for each maturity group (Figure S4A, S4B, and S4C). Grain yield, transpiration, and soil water at flowering time and biomass at the flowering time exhibited the following three interactions as significant: G_M_ × E × M (*p* < 0.02). Pairwise comparisons among these interactions revealed differences for each maturity group and planting dates in environments WW and LD, but these differences become smaller in MD and ED (Figure S4A, S4B, and S4C). In these environments, for all maturity groups, grain yield for planting dates in early May and mid-May were similar, but these differed (*p <* 0.01) from the early June planting.

### Sensitivity of initial soil moisture on LT and variability of LT in different environments

To test the effect of initial water content on the LT trait, we designed simulations and created eight scenarios of initial soil moisture (from 20% to 90%) while maintaining the LT parameter at 0.9 mm h^-1^. We conducted these simulations in Hays (Kansas), which presented the four environment classes identified in this study (Figure 2D and 5B). Nevertheless, regardless of the initial water content scenario, the model pointed out a yield increase for sorghum with LT, which is more pronounced under low soil moisture (i.e. 20% and 30%). Overall, model predictions indicated that initial soil moisture changes do not affect LT’s positive effect, although the uncertainty of these changes increased under low soil moisture (Figure 6A).

**Figure 6.**
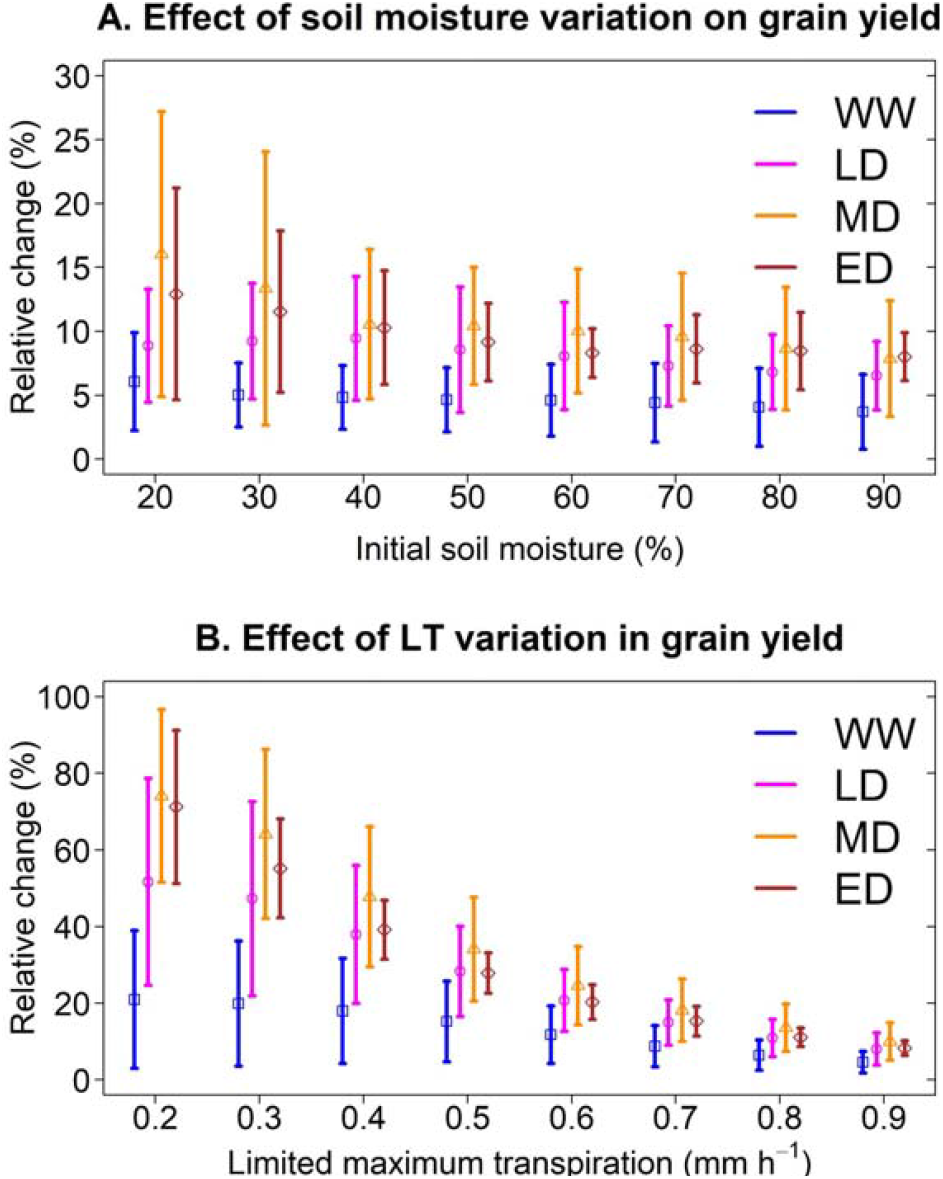
Sensitivity of a sorghum with LT trait to variation of initial conditions and genetic variability. (A) Relative change in grain yield under thresholds of initial soil moisture. The limited maximum transpiration was 0.9 mm h^-1^. (B) Relative change in grain yield under thresholds of limited maximum transpiration. Initial soil moisture was 60%. The analysis was conducted for a representative location (Hays, Kansas; 1986-2018) at the center of the sorghum belt which presented all four water stress environments. Vertical lines represent the standard deviation of each environment.

To test the hypothetical genetic variability of LT on grain yield, we created simulations for LT sorghum with a limited maximum transpiration rate from 0.2 mm h^-1^ to 0.9 mm h^-1^, while maintaining the initial soil moisture at 60%. We conducted these simulations in a central site of the sorghum belt in Kansas (Hays), representing the four water deficit patterns identified in this study. Model predictions indicated that grain yield increases in all environments when LT is lower than 0.9 mm h^-1^ (Figure 6B), with a greater benefit under drought scenarios. For instance, on average, an LT sorghum with a limited maximum transpiration rate of 0.2 mm h^-1^ increased the grain yield by 15%, 45%, 70%, and 74% in WW, LD, MD, and ED, respectively. By contrast, an LT sorghum with 0.8 mm h^-1^ increased the grain yield by 7%, 8%, 10%, and 11% in WW, LD, MD, and ED, respectively. Note that the uncertainty of predictions, represented by the standard deviation, became larger at LT lower than 0.9 mm h^-1^. In LD, the model predicted a yield increase between 25 to 79% for sorghum with an LT of 0.2 mm h^-1^. Otherwise, this increase ranged from 6 to 16% for sorghum with an LT of 0.8 mm h^-1^

## DISCUSSION

### LT for the US sorghum belt: Is it worth pursuing?

The decision to include a trait within a breeding program clearly depends on the impact of this trait on final grain yield. Breeding programs require that a candidate trait can contribute at least a 5% yield increase to be included in a breeding portfolio. Findings of this study revealed the LT trait can potentially increase grain yield from 3% to 13% in the sorghum belt in the United States (Figure 4E). Therefore, LT is a candidate trait for developing hybrids with improved water- resiliency for western regions of the sorghum belt (Figure 5B).

Although our simulation does not present a full geospatial analysis (Messina et al., 2015), our study shows results for contrasting sites across gradients of VPD and precipitation. Site-specific simulations allowed for handling detailed information on additional variables (Collins et al., 2021) in any growing period, such as soil moisture, transpiration, and biomass (Figure 5C). Otherwise, grid geospatial simulation studies rarely provide information other than yield (Guiguitant et al., 2017; Messina et al., 2015). Despite our study disregarded spatial variability on initial soil moisture, the model reproduced the observed yield (RMSE 1.1 Mg ha^-1^, Figure). Likewise, a sensitivity analysis revealed that the initial water conditions do not affect the positive impact on LT (Figure 6A).

Current climate variability (33 years) highlights the crop vulnerability (Figure 4A) in western regions characterized by the high frequency of water deficit scenarios (Figure 5B) and where the impact of LT sorghum is more significant (Figure 4G and Figure 5). Climate change scenarios, disregarded in our simulations, project a VPD increase by the end of the century (Yuan et al., 2019). Under high VPD, LT hypothetically leads to stomatal closure (Sinclair et al., 2005); similarly, rising levels of CO_2_ cause stomatal closure in C3 and C4 crops (Allen et al., 2011). However, it is unknown whether the impact of CO_2_ and LT would have a synergistic effect or if the stomatal response to CO_2_ would prevail over LT. Simulations under future scenarios would be needed to elucidate these effects. Although, a simulation study hypothesized that CO_2_ and LT can compensate for detrimental impacts of climate change in the wheat belt of Australia (Collins et al., 2021).

### Navigating G × E × M for deployment of LT

The LT trait is expected to restrict water transpiration under good soil moisture and high VPD (Sinclair et al., 2005). Therefore, this trait is undesirable for WW conditions because depriving transpiration would penalize carbon fixation and final grain yield (Vadez et al., 2014). Unexpectedly, in our study, simulation studies indicated that an LT sorghum can contribute to an increase in grain yield of 4% for WW environments (Table S3, Figure 5C). Under WW environments, predictions for wheat with the APSIM model indicated a yield increase of 0.2% (Collins et al., 2021), while predictions for maize with a simple model indicated a yield decline of -3% in the USA (mesna). Yield gains for WW environments in our study can be due to differences in the model structure. In LD environments, sorghum grain yield increased by around 9 % (Figure 5C, Table S3), which is slightly higher than predictions for wheat (2 to 7 %, Collins et al., 2021) and within the range of 0 to 24% reported for maize (Messina et al., 2015). Our results for MD (10%) and ED (9%) align with the reported yield increase for wheat which ranged between 3 to 13% (Collins et al., 2021). From a breeding perspective, LT sorghum would have a more significant impact on water stress scenarios of the sorghum belt. It is essential to identify the best combination of G_T_ × G_M_ × M in sorghum since it is planted late and across precipitation gradients (Ciampitti et al., 2019; Roozeboom and Fjell, 1998; Shroyer et al., 1998). Overall, LT sorghum increased grain yield across planting dates and maturity groups by 8%. Although specific combinations of G_M_ × M (Table S3) can maximize crop yield either for a non-LT (Figure 4C) or LT sorghum (Figure 4D).

Variety trials or multi-environment experiments have shown that, unlike full-season varieties, medium- and short-season varieties can complete their growing cycle in regions with low precipitation (Larson et al., 2021; Schnell et al., 2021) and limited growing degree days (GDD) at higher latitudes (Kukal and Irmak, 2018). This strategy has led to the selection of maturity groups that match precipitation and GDD gradients in the sorghum belt (Figures 4C and 4D). Since the impact of LT sorghum is more relevant in western regions (Figure 4E), this study suggests introgressing this trait in medium- and short-season hybrids rather than in full-season hybrids (Figure 4D, Figure 5B). Management practices need to be considered to enhance the performance of LT sorghum. Shifting planting dates can change the frequency of water deficit environments (Chenu et al., 2011) (Figure 3S) by increasing grain yield in early planting dates, especially in Kansas (Figure 4B). Higher yields in early spring resulted from the synchronization of planting dates with the onset of precipitation, which increased the frequency of WW environments (Figure S3). Likewise, simulation and field studies demonstrated yield gains of up to 11% in seasons with higher water availability (Carcedo et al., 2021; Francis et al., 1986; Zander et al., 2021)

Genetic pyramiding for drought adapted phenotypes can be explored via crop modeling (Cooper et al., 2002). A simulation study in sorghum revealed that LT is more effective than stay-green in water scarcity scenarios (Kholová et al., 2014). Higher yields in early spring suggests (Figure 4B and Figure S4D-F) a plausible interaction between early chilling tolerance and LT trait. LT increases canopy temperature (Belko et al., 2013), and temperatures beyond 38 °C can penalize carbon assimilation and plant growth (Singh et al., 2015). Then, integrating field experimentation and crop modeling for ideotyping LT with additional adaptation traits can support breeding programs when developing a trait technology for water-limited scenarios.

### Next steps for water-optimized sorghum

This simulation study has shown that LT trait can increase water productivity and benefit farmers’ economies in the sorghum belt. Nevertheless, LT is a hypothetical trait, implemented in crop models (Bates et al., 2015; Messina et al., 2015; Sinclair et al., 2017; Truong et al., 2017), and whose genetic variation is reported and limited to controlled environments (Collins et al., 2021; Devi and Reddy, 2018; Gholipoor et al., 2010; Medina et al., 2019; Vadez et al., 2015). Although variation for LT was reported in controlled environments, the repeatability of this trait has yet to be shown in sorghum-producing regions. Hence, including the LT trait in a sorghum breeding program requires validating this trait under field conditions and testing the effectiveness of phenomic approaches to discriminate this trait in large populations. Potential donors would make feasible developing bi-parental populations to determine quantitative trait loci (QTLs) controlling the phenotypic expression of this trait. Further isolating these QTL can assist in dissecting the underlying physiological and molecular mechanisms of limited transpiration, which remain enigmatic.

## ACKNOWLEDGEMENTS

This study was supported by funding from the Foundation for Food and Agriculture Research - Seeding Solution “CA18-SS-0000000094 – Bridging the Genome-to-Phenome Breeding Gap for Water-Efficient Crop Yields (G2P Bridge)” to G.P.M. and S.S.B; the Kansas Department of Agriculture “Collaborative Sorghum Investment Program Water Optimized Sorghum for Kansas” to G.P.M; and the Kansas Grain Sorghum Commission.

## AUTHOR CONTRIBUTIONS

G.P.M. and R.R. contributed to the conception and design of the work. R.R. collected experimental data, conducted simulations, data analysis, interpretation, and drafting of the article. S.S. contributed to data collection and critical revision of the article. G.M. provided technical support with the APSIM-sorghum model and critical revision of the article. All authors contributed to the final manuscript.

