## Supplemental Tables for "Crop modeling suggests limited transpiration would increase yield of sorghum across drought-prone regions of the United States"

**Supplementary information**

**Table S1. Field experiments for APSIM-sorghum testing.**

| **Location** | **Year** | **Planting date** | **Target population** | **Purpose** |
| --- | --- | --- | --- | --- |
| Colby | 1997 | 05-Jun | 25000 | model evaluation |
|  | 1998 | 02-Jun | 25000 | model evaluation |
|  | 1999 | 26-May | 25000 | model evaluation |
|  | 2000 | 01-Jun | 25000 | model evaluation |
|  | 2001 | 25-May | 25000 | model evaluation |
|  | 2002 | 30-May | 25000 | model evaluation |
|  | 2003 | 29-May | 25000 | model evaluation |
|  | 2004 | 27-May | 25000 | model evaluation |
| Garden City | 1997 | 06-Jun | 35000 | model evaluation |
|  | 1998 | 22-May | 35000 | model evaluation |
|  | 1999 | 24-May | 35000 | model evaluation |
|  | 2003 | 29-May | 35000 | model evaluation |
|  | 2004 | 01-Jun | 35000 | model evaluation |
| Hays | 1997 | 23-May | 35000 | model evaluation |
|  | 1998 | 26-May | 35000 | model evaluation |
|  | 1999 | 01-Jun | 35000 | model evaluation |
|  | 2001 | 24-May | 35000 | model evaluation |
|  | 2002 | 01-Jun | 35000 | model evaluation |
|  | 2003 | 22-May | 35000 | model evaluation |
|  | 2004 | 24-May | 35000 | model evaluation |
| Manhattan | 2007 | 05-Jun | 55000 | model calibration |

**Table S2. Cultivar parameters for a default cultivar for non-LT and LT traits. Parameters tt_endjuv_to_init (*), Tpla_prod_coef (**), and Tpla_inflection (***) for a short-season sorghum are presented in the table. Parameters for medium- and full season sorghum are indicated below the table.**

| **Parameter name** | **Unit** | **Default** | **non-LT** | **LT** |
| --- | --- | --- | --- | --- |
| tt_emerg_to_endjuv | ( °C day) | 100 | 100 | 100 |
| est_days_endjuv_to_init | () | 15 | 20 | 20 |
| pp_endjuv_to_init |  | 30 | 30 | 30 |
| tt_endjuv_to_init* | (°C day) | 138 | 138 | 138 |
| photoperiod_crit1 | (hours) | 12.3 | 0 | 0 |
| photoperiod_crit2 | (hours) | 14.6 | 13.5 | 13.5 |
| photoperiod_slope | (°C hour^-1^) | 25 | 0 | 0 |
| tt_flower_to_maturity | (°C day) | 695 | 810 | 810 |
| tt_flag_to_flower | (°C day) | 100 | 80 | 80 |
| tt_flower_to_start_grain | (°C day) | 30 | 50 | 50 |
| tt_maturity_to_ripe | (°C day) | 1 | 1 | 1 |
| Main_stem_coeff | (°C^-1^) | 2.95 | 2.95 | 2.95 |
| Tpla_prod_coef** | (°C^-1^) | 0.015 | 0.015 | 0.015 |
| Tpla_inflection*** | (°C) | 320 | 320 | 320 |
| Spla_prod_coef | (°C^-1^) | 0.007 | 0.005 | 0.005 |
| dm_per_seed | (g) | 0.0008 | 0.00099 | 0.00099 |
| plant canopy height | (mm) | 0 2000 | 0 1200 | 0 1200 |
| limited transpiration | (mm h^-1^) | - | - | 0.9 |

*	medium-season: 190, full season: 240

**	medium-season: 0.018, full season: 0.018

***	medium-season: 355.7, full season: 400.8

**Table S3. Analysis of variance for the main effects trait (G_T_), maturity group** (**G_M_), environment** (**E), and planting date (M) and its interaction on grain yield.**

| **Sour. of variation** | **df** | ***p*-val^1^** | ***p*-val^2^** | ***p*-val^3^** | ***p*-val^4^** | ***p*-val^5^** | ***p*-val^6^** |
| --- | --- | --- | --- | --- | --- | --- | --- |
| Trait (G_T_) | 1 | *** |  | *** | **** | **** | **** |
| Maturity (G_M_) | 2 | *** | *** | *** | *** | *** | *** |
| Planting date (M) | 2 | *** | *** | *** | *** | *** | *** |
| Environment (E) | 3 | *** | *** | *** | *** | *** | *** |
| G_T_ : G_M_ | 2 |  |  |  |  | *** | *** |
| G_T_ : M | 2 |  |  |  |  |  |  |
| G_T_ : E | 3 | *** | *** | *** | *** | *** | *** |
| G_M_ : M | 4 | *** |  |  |  | * | *** |
| E : M | 6 | *** | *** | *** | *** | *** | *** |
| E : G_M_ | 6 | *** | * | *** | *** | *** | *** |
| G_T_ : G_M_ : M | 6 |  |  |  |  |  |  |
| G_T_ : E : M | 6 |  |  |  |  |  |  |
| G_T_ : G_M_ : E | 6 |  |  | ** |  |  |  |
| G_M_ : E : M | 6 | * | * |  | *** | * |  |
| G_T_ : G_M_ : E : M | 12 |  |  |  |  |  |  |

^1^ Significance for grain yield, ^2^ total transpiration, ^3^ transpiration at flowering time, ^4^ soil water content at flowering, ^5^ biomass at flowering time, and ^6^ water productivity.

| **Trait (G_T_)** | **Envirom**  **(E)** | **Grain yield**  **(Mg ha^-1^)** | **Group** |
| --- | --- | --- | --- |
| non-LT | ED | 3.9 ± 0.07 | a |
| LT | ED | 4.3 ± 0.07 | b |
| non-LT | MD | 4.1 ± 0.07 | a |
| LT | MD | 4.5± 0.07 | b |
| non-LT | LD | 4.2 ± 0.08 | a |
| LT | LD | 4.6 ± 0.08 | b |
| non_LT | WW | 4.8 ± 0.08 | a |
| LT | WW | 5.0 ± 0.08 | b |

| **A** | 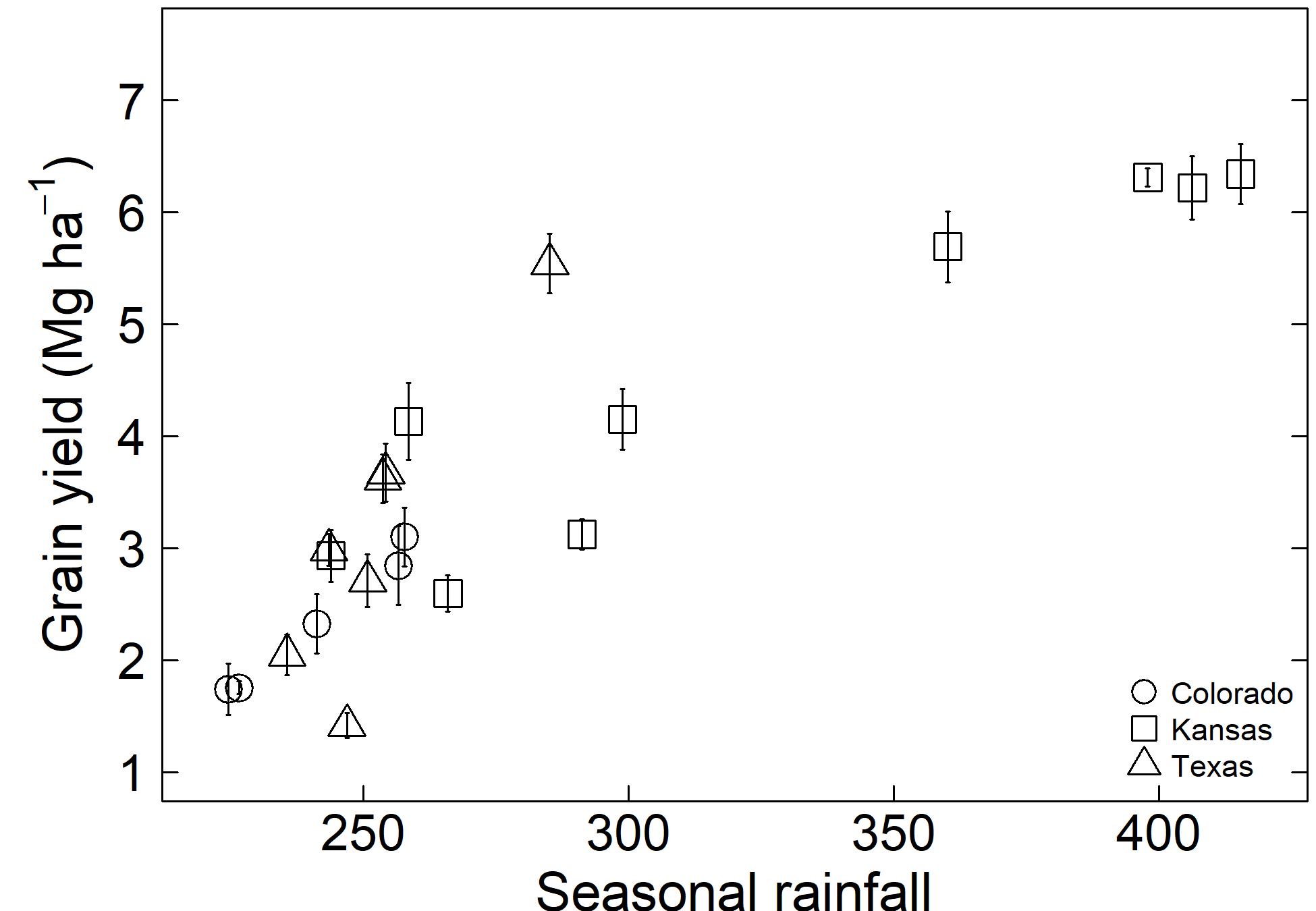 |
| --- | --- |
| **B** | 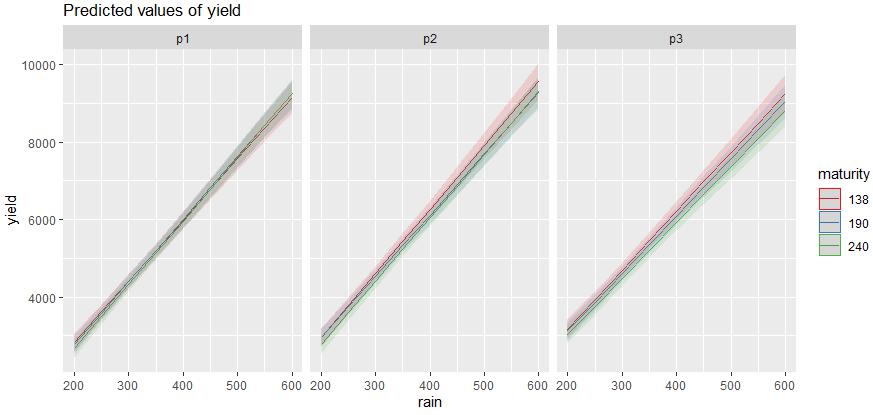 |

**Figure S1. The effect of seasonal precipitation on grain yield for a sorghum with non-LT trait.** (A) Grain yield for a simulated sorghum with non-LT trait. Simulations were conducted for three maturity groups (G_M_) on three planting dates (M). Each point represents the mean of 33 years and vertical lines indicate interannual variability (standard deviation). (B) The interaction of maturity group (G_M_) and planting date (M) across precipitation gradients for a sorghum with non-LT trait. G_M_ is represented via 138, 190 and 240 for early, medium-, and full-season sorghum, respectively. M is represented via p1, p2, and p3 for May 01, May 15 and June 01, respectively.

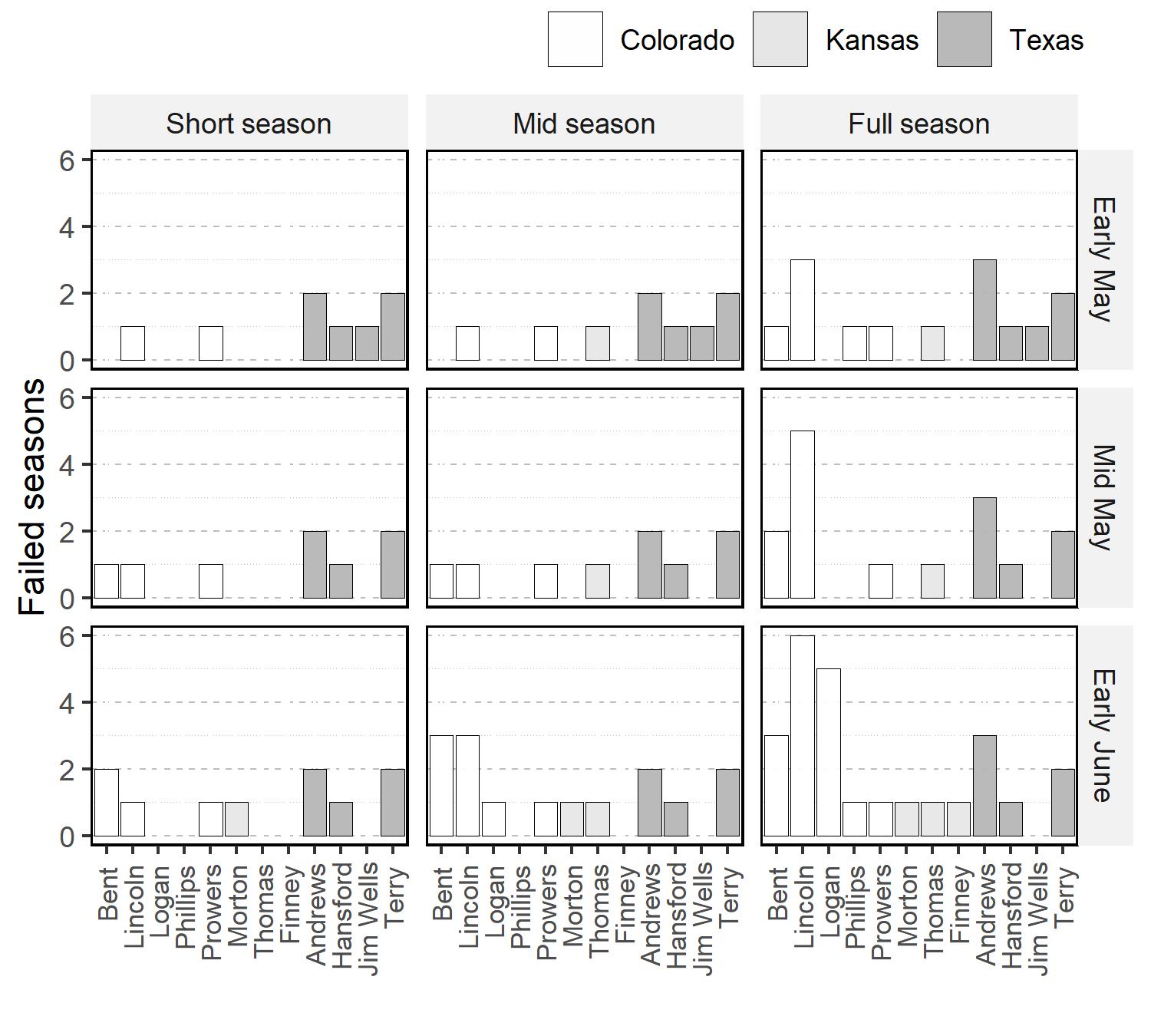

**Figure S2. Number of failed simulated seasons in the study site. A failed season indicated that the crop did not reach the vegetative or reproductive stage.**

| **A**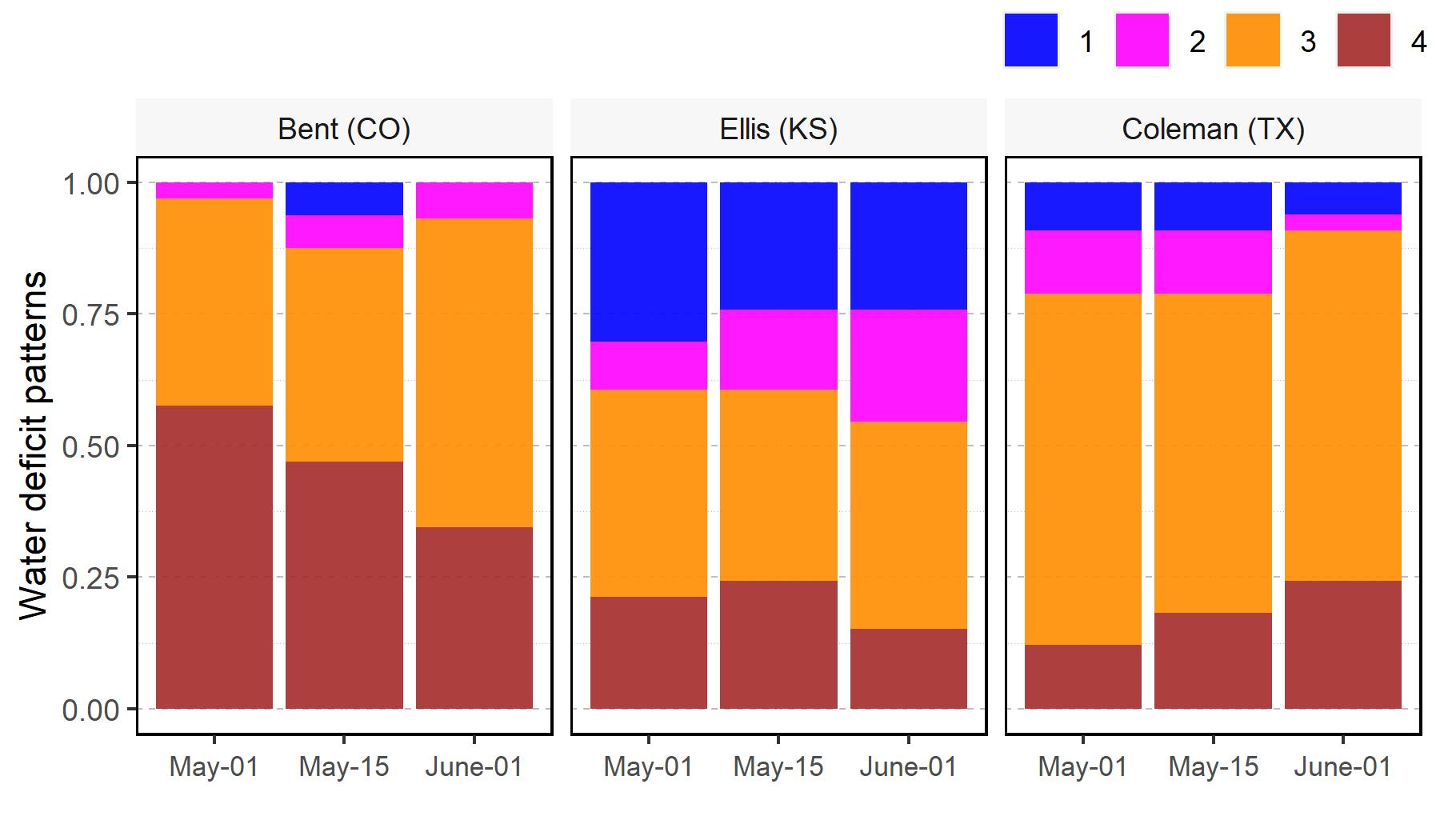 | **B**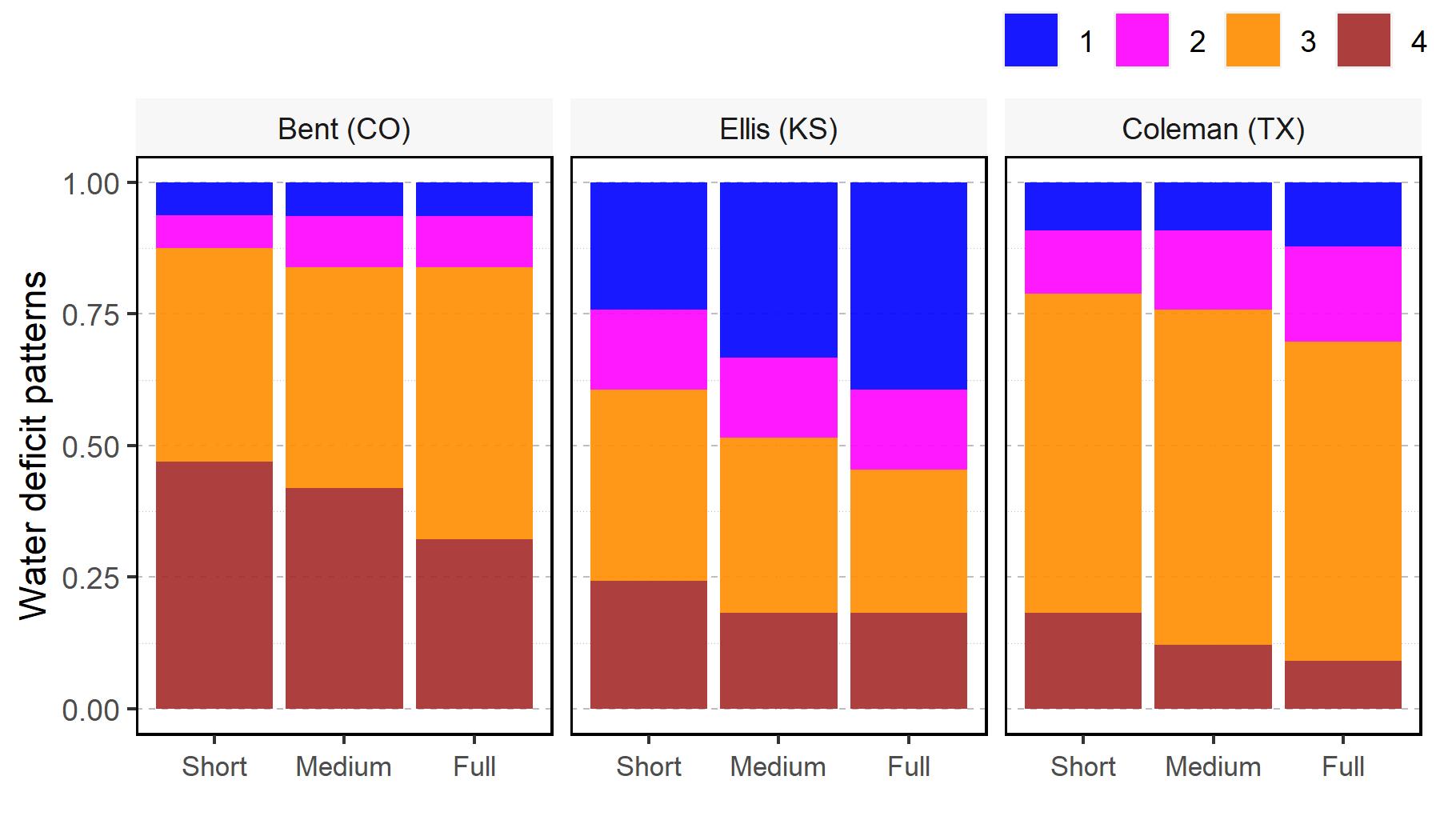 |
| --- | --- |
| **C**  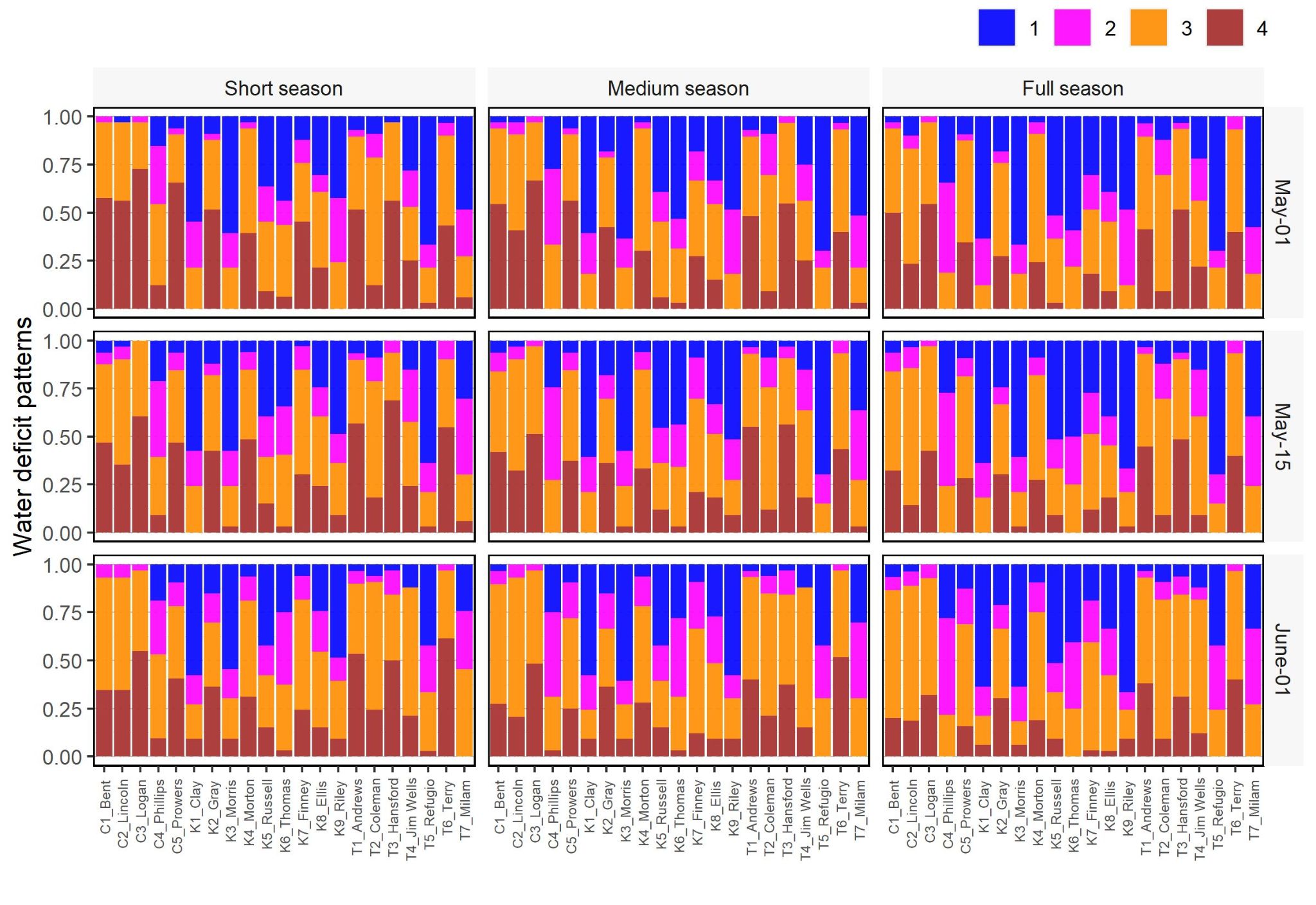 | |

**Figure S3. Frequency of water deficit scenarios in the study site.** (A) The effect of planting date on water deficit patterns for a medium season variety in Bent (CO), Ellis (KS), and Colleman (TX). (B) The effect of maturity groups on water deficit patterns for a planting date in mid-May in Bent (CO), Ellis (KS), and Colleman (TX). (C) Frequency of water deficit patterns for different maturity groups and planting dates.

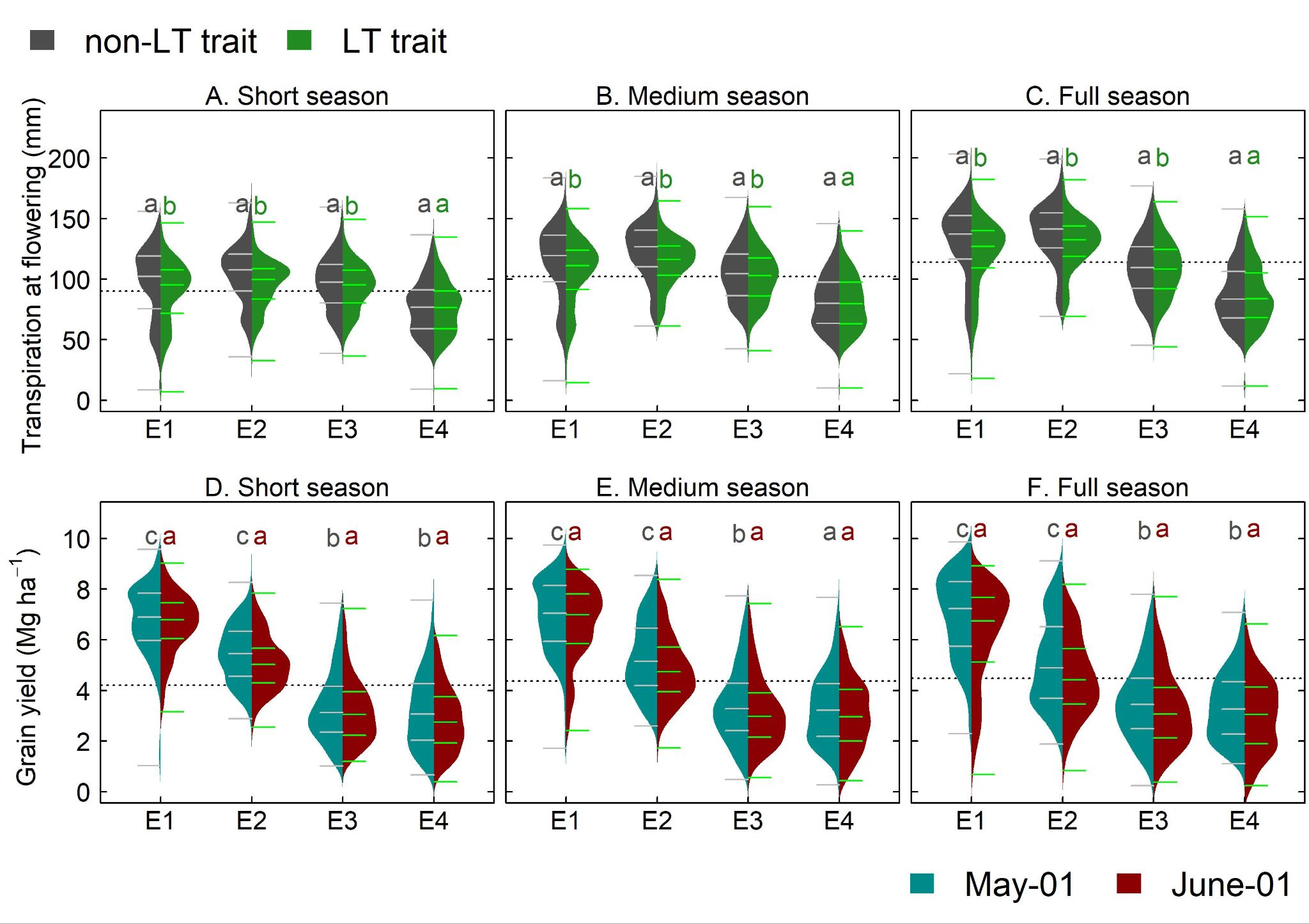

**Figure S4. Water budgets for simulated sorghum with non-LT and LT traits.** (A) soil water at flowering, (B) Cumulative transpiration at flowering, (C) Biomass at flowering, (D) total transpiration during the growing season, and (E) water productivity.

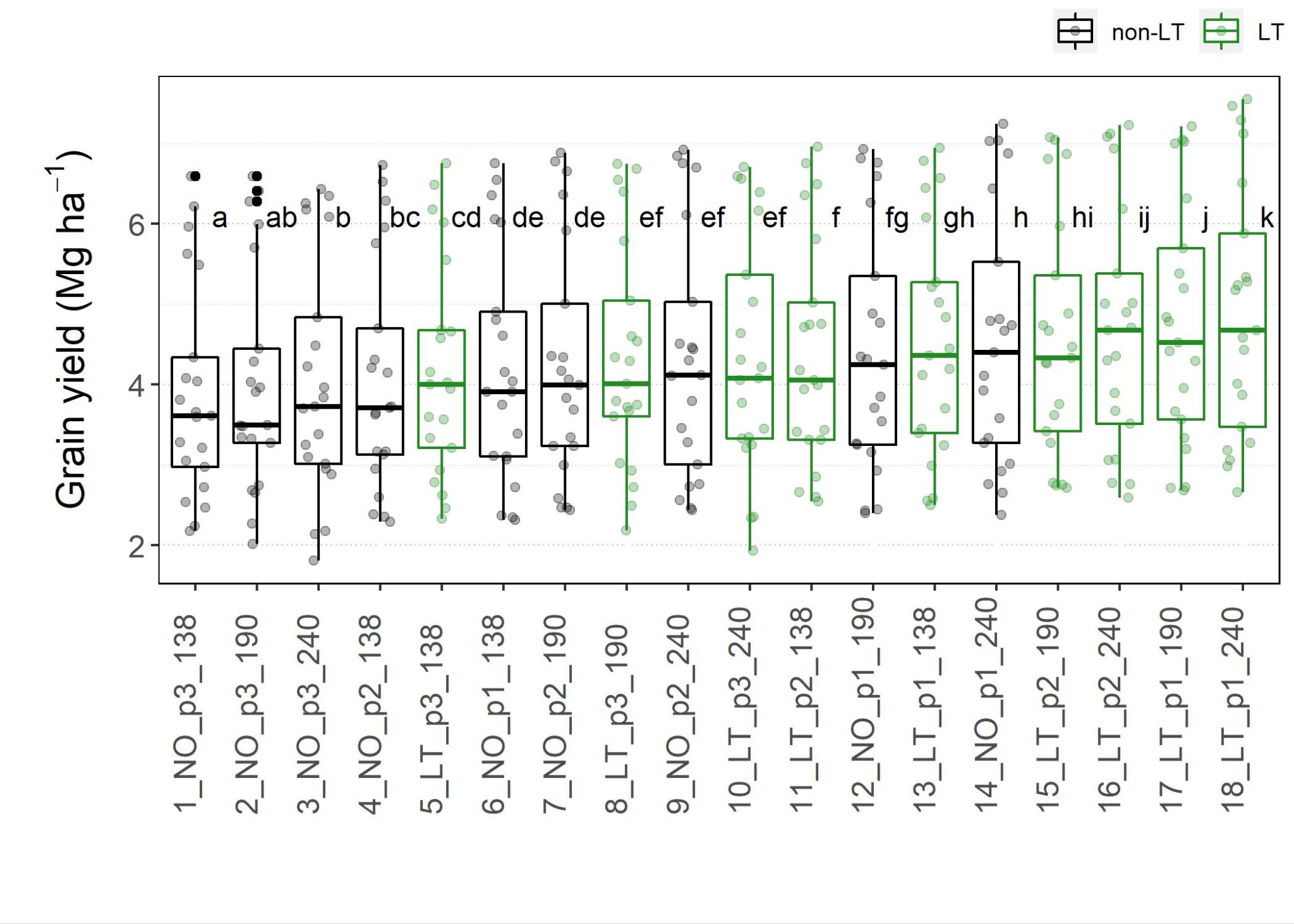

**Figure S5. Grain yield for maturity group (G_M_)** × **planting date (M)**. G_M_ is represented via 138, 190 and 240 for early, medium-, and full-season sorghum, respectively. M is represented via p1, p2, and p3 for May 01, May 15 and June 01, respectively.

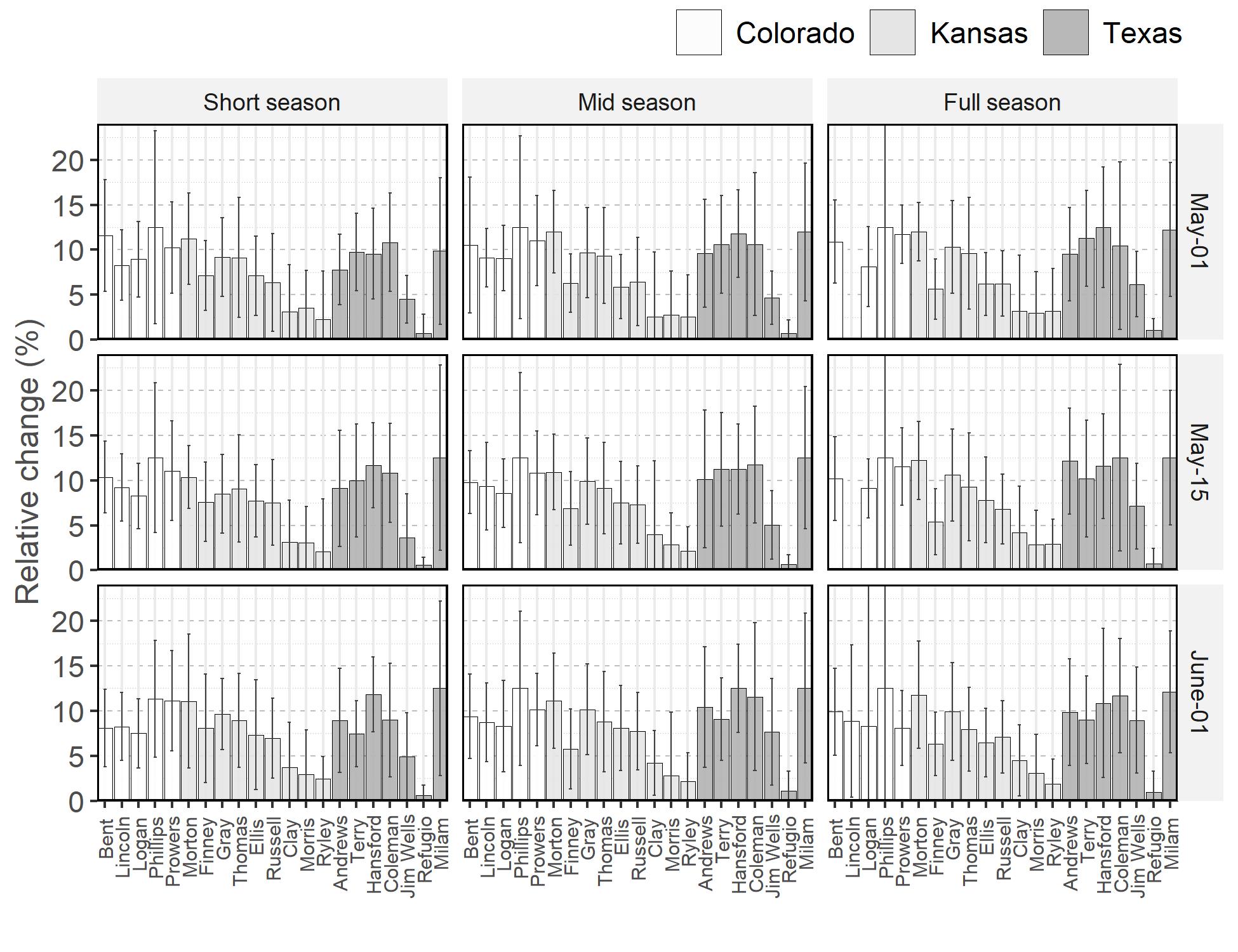

**Figure S6. Simulated mean relative change in grain yield for sorghum with LT trait.** Each barplot represents the mean from 1986 to 2018 and vertical lines indicate interannual variability (standard deviation).
